# A genome-guided atlas to the composition, activity, and β-bungarotoxin dimerization in many-banded krait venom inferred by functional venomics

**DOI:** 10.1101/2025.04.22.649985

**Authors:** Lilien Uhrig, Ignazio Avella, Lennart Schulte, Sabine Hurka, Ludwig Dersch, Johanna Eichberg, Susanne Schiffmann, Marina Henke, Ulrich Kuch, Alfredo Cabrera-Orefice, Kornelia Hardes, Andreas Vilcinskas, Maik Damm, Tim Lüddecke

## Abstract

Snakebite is a neglected tropical disease claiming ∼140,000 lives every year. One of the most medically relevant snakes of Asia is the many-banded krait (*Bungarus multicinctus*). Approximately 8% of the global human population is at risk of being envenomated by this species, able to cause fatal neurotoxicity. Here, we present a proteogenomic and functional assessment of the *B. multicinctus* venom via genome-guided bottom-up and top-down proteomics, combined with protein profiling and bioactivity assays. We report its venom profile alongside the primary structures of its toxins, revealing a relatively simple venom containing 55 components from 16 protein families. It is largely composed by three-finger toxins and phospholipase A_2_, besides acetylcholinesterase and snake venom metalloprotease. Top-down data unveiled the diversity of the highly lethal β-bungarotoxins and allowed us to infer the complex dimerization patterning of these multi-domain neurotoxins. Our functional analysis revealed that *B. multicinctus* venom exerts potent phospholipase A_2_ and acetylcholinesterase activities, while protease activity and effects on cell viability and release of second messengers were virtually absent. This suggests, that *B. multicinctus* venom causes its devastating neurotoxic symptoms due to a heavy reliance on phospholipase A_2_ and acetylcholinesterase, but without directly impairing neuron viability or via interference of second messenger release. Antibacterial and antiviral screens further revealed activity against some pathogenic microbes that warrant further translational investigations. A comparison to previously published venom proteomes of *B. multicinctus* and its congeners suggests, that intraspecific venom variation occurs more widely in kraits than previously acknowledged and deserves higher attention. Overall, our investigation provides pivotal new insights into the biochemistry and pathophysiology of one of Earth’s most lethal snakes and represents an important resource to inform future proteogenomic and functional studies in krait venom and beyond.

## 2 Introduction

Snakebite is a global health burden that primarily affects economically disadvantaged communities in the global south [1–4]. Annually, an estimate of 5.4 million snakebites cause approximately 140,000 deaths and 400,000 cases of long-term disabilities [4]. However, cases occur primarily in rural settings, thus only a small fraction is reported and available metrics are likely a severe underestimation of the true global burden of snakebite [5,6]. In recognition of its medical importance, the World Health Organization (WHO) in 2017 declared snakebite envenoming a neglected tropical disease and set a strategic goal to halve its mortality and morbidity by 2030 [7]. To achieve this, a series of strategies has been proposed, of which the improvement in availability and effectiveness of treatment measures is particularly important [8]. However, its achievement is intrinsically linked to a deep compositional and mechanistic understanding of the venom toxins, which feature the functional drivers of envenoming symptoms [9,10].

Snake venoms are complex cocktails dominated by toxic enzymes and peptides [11,12]. Their molecular diversity is increased due to its adaptive potential, leading to large extents of venom variation across all taxonomic levels [13–16]. In light of this complexity, the efficient characterization of venom profiles has long been a challenging task and many species remain under- or even unstudied thus far [17]. Recently, the emergence of a novel field, referred to as “modern venomics”, provided the means to rapidly resolve the composition and functionality of venom cocktails from across the animal kingdom [9,10]. Especially, the increased implementation of high-quality genomic data creates promising new avenues to study snake venom [18]. This is due to i) the opportunity to infer insights on venom components at the nucleotide-level scale when genomes are utilized as species-specific databases for venom proteomics, ii) to reduced error rates in comparison to transcriptomes sequenced de novo and iii) it further allows the study of evolutionary processes by inference of genomic localization and synteny [18–20]. However, despite its potential, genomic data is scarcely applied in snake venom research.

One of these highly medically important, yet understudied species is the many-banded krait (*Bungarus multicinctus*). This large (up to 180 cm body length) species belongs to the Elapidae family, which includes many of the most medically relevant snakes of the world, and is characteristically colored due to its prominent black-white banding pattern [21–23], Figure 1. The 18 species of kraits currently recognized are distributed across Asia. *Bungarus multicinctus*, specifically, can be found in South China (including Hong Kong, Hainan), Laos, Myanmar, North Vietnam (Hoa Binh), Taiwan and Thailand [24,25], Figure 1. In these regions, it occurs in a wide range of habitats such as lowland wooded areas, bamboo thickets and rice paddies, as well as villages and suburbs of larger cities [26,27]. Based on this wide distribution across one of the more population-rich areas of Asia, the WHO estimates that more than 650 million people live within the range of *B. multicinctus*. This renders a theoretical 8% of the world population being at risk of envenomation by this species [23]. Similar to other kraits, it possesses a potent neurotoxic venom known to cause lethal respiratory failure when untreated [28,29]. Victims often get bitten at night while sleeping, and tend to remain oblivious as the bite is painless and symptoms can be absent for up to 6 hours [30–33]. Because of this, krait bites may go undetected and thus remain untreated, leading to fatalities. These dreadful, yet common, cases earned kraits their nickname “silent killers” [34]. Although *B. multicinctus* is generally non-aggressive, bites occur frequently. Due to their high mortality rate and large human population at risk, this species is classified as being of “highest medical importance” and is considered one of the most dangerous snakes of Asia [23].

**Figure 1:**
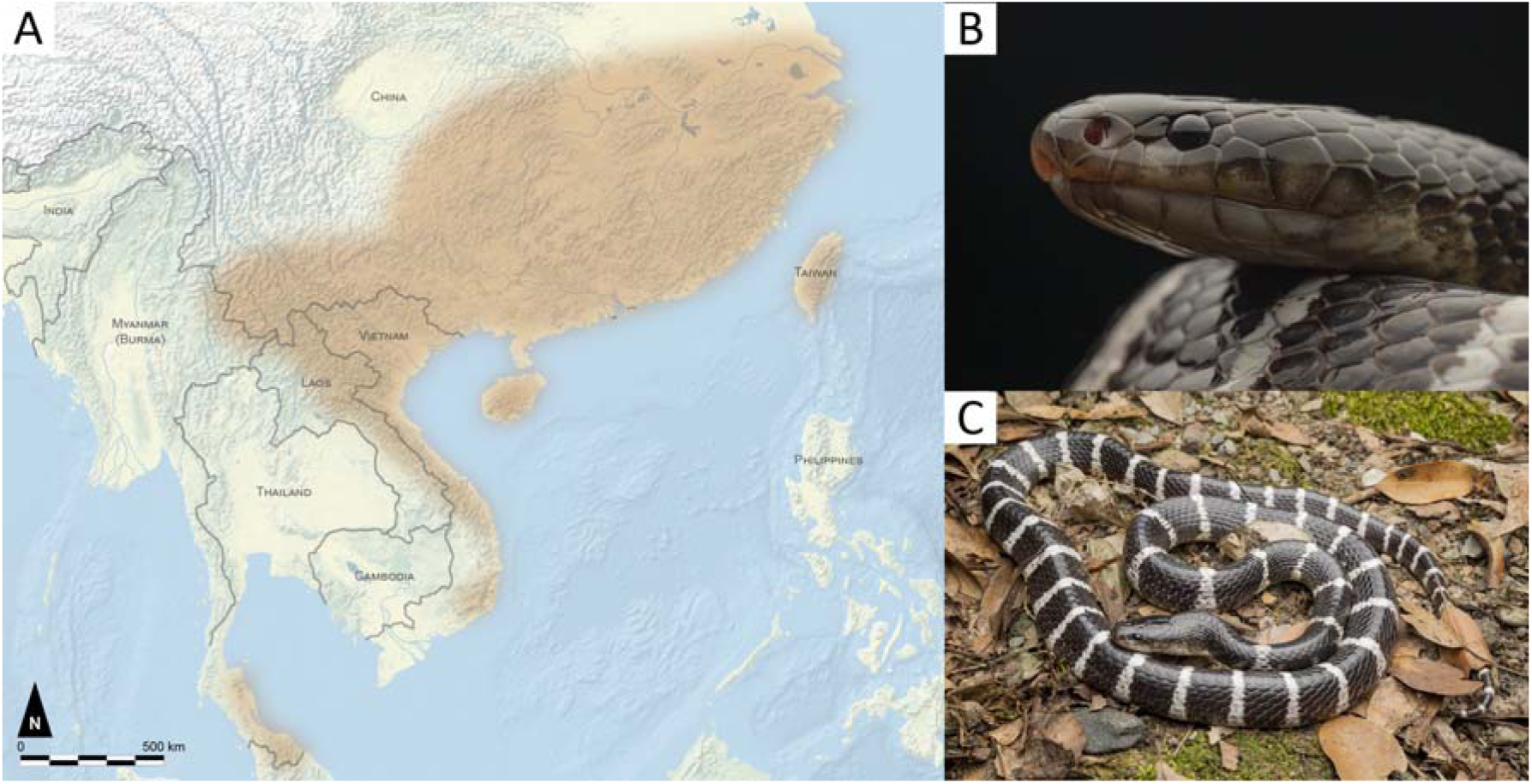
Distribution range and appearance of the many-banded krait (*Bungarus multicinctus*). (A) Distribution range of *B. multicinctus* (brown) [24]. (B) Close-up and (C) full-body photograph of an adult specimen from Taiwan. Photo credits: Kai Kolodziej.

Based on this medical relevance, a large set of previous studies investigated the venom of *B. multicinctus* for its composition and function. These works revealed a venom rich in three-finger toxins (3FTx) and phospholipases A_2_ (PLA_2_) [35–37] in agreement to the neurotoxic symptoms caused [38]. In addition, recently published reference genomes for *B. multicinctus* provided a theoretical basis to apply genome-guided modern venomics to provide the highest possible resolution when deciphering its venom profile [39–41]. Nevertheless, despite this considerable potential to investigate this highly important species, little effort has been made to utilize the available genomic repositories and hence many important questions revolving around many-banded krait venom biology and chemistry remain unanswered.

Here, we compensate for this recalcitrant research gap and utilize a recently sequenced *B. multicinctus* genome together with bottom-up, and top-down proteomics as well as SDS-PAGE and RP-HPLC based profiling to provide the most detailed available atlas to the species toxin arsenal and their primary structures. This proteogenomics-guided inventory is supplemented by one of the largest bioactivity profiling thus far carried out for krait venoms to set the identified components into functional perspective. We particularly explore the *B. multicinctus* venom for its ability to exert activities reminiscent to neurotoxicity, such as PLA_2_ and acetylcholinesterase (AChE) activity, and investigate its effects across several cell types and on the release of second messengers. Our study is therefore offering a novel compositional and functional perspective upon *B. multicinctus* venom, helping to understand and treat the symptoms caused by it.

## 3 Material and Methods

### 3.1 Venom samples

Venom from nine adult *Bungarus multicinctus* specimens of unknown sex was collected in Taiwan in 2002. The samples were pooled, lyophilized and stored at 6 °C until analysis. Legal ID from Taiwan that allowed the collection?

### 3.2 Venom Profiling

#### 3.2.1 SDS-PAGE

One-dimensional SDS-PAGE profiling was carried out, using 3 µg of redissolved *B. multicinctus* venom in double-distilled water (ddH_2_O). Samples were equally mixed (v/v) with non-reducing or reducing (5% (v/v) 2-mercaptoethanol) 2 x Laemmli buffer and denatured at 95 °C for 5 min. The venom was loaded to a 4–20 % Mini-PROTEAN® TGX™ Precast Protein Gels (Bio-Rad) and separated at 120 V for 60 min in a Mini-PROTEAN® Tetra Vertical Electrophoresis Cell (Bio-Rad). The gel was stained (ROTI®Blue quick solution; Carl Roth) and destained with acetic acid (10%(v/v)), methanol (50% (v/v)). Gels were digitalized with the Molecular Imager® Gel Doc™ XR+ System (Bio-Rad) with Image Lab Software (6.1.2).

#### 3.2.2 RP-HPLC

For the RP-HPLC venom profile, *B. multicinctus* venom was dissolved to a final concentration of 10 mg/ml in 20 µl aqueous 5% (v/v) acetonitrile (ACN) with 1% (v/v) formic acid (HFo) and centrifuged for 10 min at 12,000 × g. The supernatant was measured on a reversed-phase Supelco Discovery BIO wide Pore C18-3 (4.6 × 150 mm, 3 µm particle size) column operated at 40 °C by a HPLC Agilent 1100 (Agilent Technologies, Waldbronn, Germany) chromatography system. The following gradient with ultrapure water with 0.1% (v/v) HFo (solvent A) and ACN with 0.1% (v/v) HFo (solvent B) was used at 1 ml/min, with a linear gradient between the time points, given at min (B%): 0–5 (5% const.), 5–65 (5 to 45%), 65–75 (45 to 70%), 75–80 (95% const.), and 10 min re-equilibration at 5% B. The chromatography runs were observed by a diode array detector (DAD) at λ = 214 nm detection wavelength (360 nm reference). A previous blank run (aqueous 5% (v/v) ACN with 1% (v/v) HFo, centrifuged for 10 min at 12,000 × g) was measured under identical parameters and subtracted from the venom profile.

#### 3.2.3. Bottom-Up Proteomics

To infer the protein composition of our *B. multicinctus* venom sample, we applied a modified version of a bottom-up shotgun proteomic workflow previously established for animal venom analysis [42–44]. Lyophilized venom samples were reconstituted in 1 ml of 6 M urea, 50 mM Tris/HCl pH 8.0 to ensure complete denaturation. Aliquots containing 100 µg of protein were transferred to Protein LoBind tubes (Eppendorf), and 10 mM TCEP was added to reduce disulfide bonds (incubation for 30 minutes at 50 °C). Free thiols were then alkylated with 50 mM chloroacetamide (incubation for 30 minutes at 22 °C in the dark).

The samples were diluted 1:10 with 50 mM Tris/HCl pH 8.0, 5% ACN to reduce the urea concentration to less than 1 M. Next, 1 µg of Gold Trypsin (Promega) was added at a 1:100 protease-to-protein ratio, and the venom was digested overnight at 37°C. The digestion was halted by adding trifluoroacetic acid to a final concentration of 1%. The samples were then cleaned up using C18-Chromabond columns. Peptides were eluted, dried using a vacuum concentrator (Eppendorf), and re-dissolved in 30 μl of 5% ACN, 0.1% (v/v) HFo. Peptide concentrations were determined using the Pierce colorimetric kit (Thermo Fisher Scientific), and samples were stored at −20°C until mass spectrometry (MS) analysis. Before MS measurement, peptides were separated chromatographically using a Thermo Fisher Scientific UltiMate 3000 RSLCnano system. 1 µg of peptides were loaded onto a PepMap Neo Trap column for concentration and desalting, followed by separation on a 25 cm Aurora Ultimate column (IonOpticks). The analytical column was operated at 60°C in an external oven. Peptides were eluted at 300 nl/min with a linear gradient (solvent B: ACN, 0.1% HFo) between the time points, given at min (B%): 0–80 (5 to 40%), 80–90 (40 to 90%), 75–80 (95% const.), and the column was then washed with 90% acetonitrile, 0.1% (v/v) HFo and re-equilibrated with 2% ACN, 0.1% (v/v) HFo for at least 10 minutes. MS analysis of the peptides was performed on an Orbitrap Eclipse Tribrid MS (Thermo Fisher Scientific, MA, USA) using positive ionization via electrospray at 2.2 kV and a source temperature of 275 °C. MS scans were conducted in data-dependent acquisition mode. Detailed parameters are provided in Table S15.

Mass spectrometry data was analyzed by PEAKS Studio v12.0 (Bioinformatics Solution Inc.) using a previously sequenced *B. multicinctus* genome as species-specific database for peptide-searches (see 3.2.5 Genome Annotation). In PEAKS, the DDA RAW data were analyzed by De Novo, DB Search, PEAKS PTM and SPIDER; with mass error tolerances (parent: 15.0 ppm; fragment: 0.5 Da), semi-specific trypsin digest (max missed cleavages: 3) and carbamidomethylation as fixed beside several variable modifications. A list of detailed parameters is provided in Table S10. The filtered hit list with unique amino acid sequences for at least two unique peptides and a PEAKS −10LgP >30 is provided in Table S11.

To provide information on toxin quantities, the relative abundances were estimated as Normalized Spectral Abundance Factors (NSAF) and calculated as described previously [44]. Briefly, the spectral counts (SpC), are used as a substitution for protein abundance [45,46]. The NSAF of each protein was calculated, as presented below:

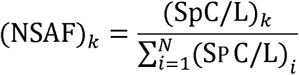

To calculate the NSAF for a protein k its SpC is normalized per its amino acid sequence length (L), against the sum of all SpC/L for N proteins [47,48]. A detailed list of all identified venom components, additional information on the calculated relative abundances, as well as a detailed list of all identified venom components is presented in Table S9.

#### 3.2.4 Top-Down Proteomics

The used top-down workflow was adapted from a previously published protocol [49] with the following changes: 200 µg venom were used for two aliquots (reduced and non-reduced) of 50 µl each. The blank consists of citrate buffer (30 µl, 100 mM, pH 3), ultra-pure water (10 µl) and 1% (v/v) HFo (10 µl).

First, 5 µl were injected into reversed-phase ACQUITY UPLC Peptide BEH C18 (300 Å, 1.7 µm, 2.1 mm × 100 mm) column with the appropriate pre-column (ACQUITY UPLC Peptide BEH C18 VanGuard Pre-column, 300Å, 1.7 µm, 2.1 mm × 5 mm) with an Agilent 1290 Infinity LC system (Agilent Technologies) coupled to an ESI QTOF maXis II mass spectrometer (Bruker Daltonics). The following gradient with ultrapure water with 0.1% (v/v) HFo (solvent A) and ACN with 0.1% (v/v) HFo (solvent B) was used at 0.6 ml/min, with a linear gradient between the time points, given at min (B%): 0–0.3 (5% const.), 0.3–18 (5 to 95%), 18–18.1 (95 to 100%), 18.1– 22.5 (100% const.), and 2.5 min re-equilibration at 5%. Data was acquired in the scan range 50–2000 m/z. MS/MS analysis was carried out for the top five most intense ions (30 s exclusion list) selected for collisional induced dissociation (CID) using N_2_.

The data analysis was adapted from a published protocol [49]. In short, the RAW files were converted to the mzML format (vendor msLevel = 1−; −; removeExtra zeroSamples = 1−)[50], further converted to the MSALIGN format using TopFD [51] and run by TopPIC [52]. To confirm the bottom-up results, identified entries from our PEAKS analysis with a maximum of one single unknown post translational modifications (PTM) were combined with all Kunitz-type serine-protease inhibitors (KUN), 3FTx and PLA_2_ sequences from the underlying original genome assembly publication [39] and UniProtKB/TrEMBL entries based on the search pattern “bungarotoxin AND (organism_id:8616)” [53]. The database was cleaned of duplicate entries. A single unknown PTM was allowed. A list of detailed parameters is provided in Table S12. For intact mass profiling (IMP) of the non-reduced and reduced top-down files, data were manual screened in the Compass DataAnalysis v5.3 (Bruker Daltonics) for an overview of abundant masses and disulfide bridge counting. A list of raw data results is provided in Table S14.

#### 3.2.5 Genome Annotation

The sequences of proteins identified via the different proteomics workflows were retrieved from the used *B. multicinctus* genome [39] and further annotated with DIAMOND v2.1.10 (sensitivity mode: ultra; max target seqs: 0/all; E-value: 1e-3) [54] against the UniProtKB 2024_06 [53] based databases Swiss-Prot, TrEMBL (serpentes, taxonomy ID: 8570), VenomZone [55] and Tox-Prot [56](all downloaded on January 26, 2025). In addition, we calculated query and subject coverage and sequence similarity based on the BLOSUM62 matrix [57] within BioPython v1.83 [58]. The results were arranged in descending order by similarity, query and subject coverage for each candidate, with the top DIAMOND hit selected for further analysis. Peptide sequences were also screened for signal peptides using SignapP v6.0h (mode: slow_sequential; organism: eukarya) [59] and annotated with InterProScan v5.72-103.0 [60]. With this information we manually assigned the putative venom components to their respective peptide families.

#### 3.2.6 Data Accessibility

Following the FAIR principle, *B. multicinctus* genome assembly data from project CNP0002662 at the China National GeneBank Database (CNGBdb) and the annotation file could be reused for our study [39]. All newly generated proteomics data have been deposited in the Zenodo repository (https://zenodo.org) under the project name “DATASET - Mass Spectrometry - Snake venom proteogenomics of *Bungarus multicinctus* from Taiwan” with the data set identifier “15118450” [61].

### 3.3 Biological activity assays

#### 3.3.1 Phospholipase A_2_ Activity Assay

The Phospholipase Activity of the *Bungarus multicinctus* venom was measured using the EnzChek™ Phospholipase A_2_ Assay Kit (Invitrogen, E10217) for 96-well plates, according to the manufacturer’s instructions. Briefly, venom was used to test final concentration of 50, 25, 12.5, 6.25 and 3.125 µg/ml in the PLA_2_ reaction buffer in a round-bottom 96-well plate. Furthermore, a positive control with isolated PLA_2_ from honey bee venom and a negative control with only reaction buffer were prepared. The fluorescence resonance energy transfer (FRET) measurements were performed 5 minutes after mixing the samples with an emission (E) intensity ratio at 515/575 nm with excitation at ∼460 nm in a microplate reader. The measured values were averaged and normalized against the negative (0%) and positive control (100%). The significance of the results was determined using a student’s t-test.

#### 3.3.2 Acetylcholinesterase Activity Assay

The Acetylcholinesterase (AChE) Activity of the venom was analyzed with the Acetylcholinesterase Activity Assay Kit (Sigma Aldrich, MAK119) according to the manufacturer’s instructions. In short, the venom was tested at final concentrations of 5, 2.5 and 1.25 µg/ml, against a negative (neg., ddH_2_O) and calibrator (equivalent to 200 units/l). Then the blank (ddH_2_O) and the calibrator (equivalent to 200 U/L) were prepared. The absorbance (A) was measured at λ = 412 nm with a Synergy H4 microplate reader (BioTek). The initial and final measurement were taken after 2 and 10 minutes of incubation. The AChE Activity was calculated as follows:

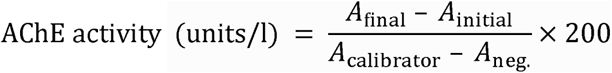

#### 3.3.3 Protease Activity Assay

To analyze the protease activity of the venoms, the “Protease Activity Kit” from Calbiochem was used, using a Fluorescein thiocarbamoyl (FTC)-casein according to the manufacturer’s instructions. In short, *Bungarus multicinctus* venom were tested at final concentrations of 50, 100 and 200 µg/ml. In a round bottom 96-well plate, 25 µl FTC casein (0.6% FTC-casein in 50 mM Tris-HCl, pH 7.3) and 25 µl incubation buffer were added. Then, in triplicates, 10 µl of ddH_2_O (negative control), 10 µl of trypsin (positive control) or 10 µl of the respective venom predilutions were added. Finally, the plate was covered with an air-permeable foil and incubated in a zip-lock bag with sheets of wet paper towel inside at 37 °C and 120 rpm. After 2 h, the reaction was stopped by adding 120 µl of a trichloroacetic acid (TCA) solution and incubated for further 20 min under the same conditions.

The plate was then centrifuged at 500 x g and 4° C for 20 min to pellet the undivided precipitated dyes. Subsequently, 40 µl of the supernatant was removed, transferred to a flat bottom 96 well plate and mixed with 160 µl assay buffer. The FTC peptides were quantified by measuring the absorbance at λ = 492 nm in a Synergy H4 microplate reader (BioTek). The plate was shaken for 20 s at 237 rpm to obtain a homogeneous solution, then the measurements were performed in triplicates. The color intensity produced is directly proportional to the protease activity in the sample. The measured values were averaged and normalized against the negative (0%) and positive control (100%). The significance of the results was determined using a student’s t-test.

#### 3.3.4 Cell Viability Assay

To investigate the cytotoxicity of *B. multicinctus* venom, cell viability assays were performed using the Orangu™ assay (Cell Guidance Systems) on the macrophage cell line RAW 264.7, peripheral blood mononuclear cells (PBMCs), human embryonic kidney cells (HEK 293T) and neuroblastoma cells SH-SY5Y as described by [62]. 2 × 10^4^ HEK293T, SH-SY5Y, or RAW246.7 cells were seeded in 96-well plates for 24 h. The cells were treated with different concentrations of *B. multicinctus* venom or ddH_2_O as a control for 24 h. 1 × 10^5^ PBMCs were seeded in 96 well plates and treated with different concentrations of *B. multicinctus* venom or ddH_2_O as a control for 24 h. 10 μl of Orangu™ cell counting solution was added, followed by 60 min (HEK293T, SH-SY5Y, RAW246.7) or 2h (PBMCs) of incubation. Absorbance was measured at λ = 450 nm with a reference wavelength of λ = 650 nm using an EnSpire 2300 Multimode Plate Reader (Perkin Elmer). Celecoxib (100 μM) and ddH_2_O were applied as positive control or negative control respectively. Measurements were normalized to the negative control (ddH_2_O), which was set to 100% cell viability.

#### 3.5.5 Antibacterial Assay

To obtain a broad screen for antibacterial activity a panel of seven bacterial strains was carried out as previously described by Hurka et al., 2022. *Escherichia coli* (DSMZ 102053), two *Pseudomonas aeruginosa* strains (DSMZ 1117 and DSMZ 50071), *Listeria monocytogenes* (DSMZ 20600), *Micrococcus luteus* (DSMZ 20030), *Staphylococcus aureus* (DSMZ 2569) and *Staphylococcus epidermidis* (DSMZ 28319) were tested against a dilution series (2 mg/ml, 1 mg/ml, 0.5 mg/ml, 0.25 µg/ml and 0.125 mg/ml) of *B. multicinctus* venom. The experiment was carried out in 96-multiwell plates with a final volume of 100 µl per well (50 µl of bacterial suspension; 50 µl of diluted venom), followed by OD600 measurements in triplicates using a BioTek Eon microplate reader 18 h after venom exposure. The bacteria growth was normalized to the solvent control cultures (100%) and antibiotic control with genatmicin (Sigma-Aldrich, Taufkirchen, Germany, 0%). The data was analyzed and visualized in GraphPad Prism v10.0.2.

#### 3.3.6 Antiviral Activity Assay

The antiviral activity Assay was carried out as previously described [63]. Three strains of influenza virus: B/Massachusetts/71 (Massachusetts/B), B/Malaysia/2506/2004 (Malaysia/B), and A/Hamburg/05/2009 (H1N1pdm) were tested against four different venom concentrations (12.5 µg/ml, 25 µg/ml, 50 µg/ml and 100 µg/ml) as well as 10 mM aprotinin (Carl Roth, A162) as a positive control and an untreated negative control. Viruses were propagated in MDCK-II cells in an infection medium containing DMEM GlutaMAX supplemented with 1% penicillin/streptomycin, 0.2% bovine serum albumin (BSA, Thermo Fisher Scientific) and 1Cµg/mL bovine N-tosyl-l-phenylalanine chloromethyl ketone (TPCK)-treated trypsin (Thermo Fisher Scientific). The cells were then seeded to 90% confluence in the infection medium excluding trypsin and inoculated with A/Hamburg/05/2009 (H1N1pdm), B/Malaysia/2506/2004 (Malaysia/B) at a multiplicity of infection of 1 (MOI = 1) and B/Massachusetts/71 (Massachusetts/B) at MOI = 0.1. After 1 h of incubation the cells were washed twice and then treated with the dissolved venom or aprotinin in the infection medium. Cell viability was assessed 48 h later by measuring the ATP content using the CellTiter-Glo Luminescent Cell Viability assay (Promega) according to the manufacturer’s instructions, measuring luminescence in black 96-well plates using a Synergy H4 microplate reader (BioTek). Luminescence values normalizing to the aprotinin control (100%) and blank (virus-treated cells, 0%). For calculating means and standard deviations triplicate measurements were used.

#### 3.3.7 Assessment of intracellular Ca^2+^ levels

The assessment of Ca^2+^ release was performed by labelling cells with the calcium fluorescence probe Fluo-8-AM, as described by [62]. After 2 × 10^4^ HEK 293T or 4 × 10^4^ SH-SY5Y cells were plated in 96-well poly-d-lysine-coated plates and incubated at 37 °C for 24 h the cells were treated with 4.19 μg/ml Fluo-8-AM in Hanks’ balanced salt solution (HBSS) for 1 h at 37 °C. The Fluo-8/HBSS medium was replaced with 100 μl of HBSS. For imaging an ImageXpress Micro confocal high-content imaging system (Molecular Devices, San Jose, CA, USA) was used capturing five pictures over five seconds. In the inhibition assay 5 μM ionomycin was added to the venom-treated samples after 30 min, and images were every second for 20 s. For the induction assay, cells were exposed to vehicle (negative control), *B. multicintus* venom, or 5 μM ionomycin (positive control, Sigma-Aldrich), and images were taken as above. The data was analyzed utilizing MetaXpress software. Cells exhibiting a fluorescence intensity above the threshold, based on pre-treatment cells, were quantified. For the induction assay, the number of cells surpassing the threshold in venom-treated samples was compared with the untreated samples. In the inhibition assay, venom-treated samples were compared with cells treated with ionomycin. Results of the induction and inhibition assay were normalized against the positive control (100%) and against the negative control (0%).

#### 3.3.8 Assessment of NO levels

The assessment of nitric oxide (NO) levels of *Bungarus multicinctus* venom was performed following [62]. In short, 2 × 10^4^ RAW264.7 macrophages per well were seeded in a 96-well plate and incubated for 24 h at 37°C. The venom was added at final concentrations of 0.25, 2.5 and 25 µg/ml as well as vehicle (negative control) or 100 ng/ml lipopolysaccharide (LPS, positive control) or to induce NO synthesis. For 30 min the cells were pre-treated with venom or vehicle before being incubated with 100 ng/ml LPS to investigate the inhibition of NO synthesis. Supernatants were collected and stored after 24 h at –80°C. Using different concentrations of sodium nitrite (0–50 μM) a standard curve was generated. Then 80 μl cell supernatant or standard were added to a 96-well microplate and incubated with 20 μl sulfanilamide solution (40 mg/ml in 1 M HCl) and 20 μl naphthylenediamine solution (60 mg/ml N-(1-naphthyl)ethylenediamine dihydrochloride in water) for 15 min. The absorbance was measured at λ = 540 nm using an EnSpire Plate Reader (Perkin Elmer). Results of the induction and inhibition assay were normalized against the positive control (100%) and against the negative control (0%).

## 4 Results

### 4.1 Composition of the *Bungarus multicinctus* venom proteome

For an initial exploration of the molecular complexity of *Bungarus multicinctus* venom we performed venom-profiling by one-dimensional SDS-PAGE and RP-HPLC (Figure 2). Under non-reducing conditions, two conspicuous band areas can be observed at 20 kDa and in the range of 10 to 15 kDa, whereas under reducing conditions we identified two bands at ∼15 kDa and ∼10 kDa. The apparent masses are in line with putative toxin families prominent in *Bungarus* venom, including three-finger-toxins (3FTx, 6-8 kDa) such as alpha-bungarotoxin (α-BT), kappa-bungarotoxin (κ-BTx), gamma-bungarotoxin (γ-BTx), muscarinic toxins (MT). They are further reminiscent to phospholipase A_2_ (PLA_2_, 13-14 kDa) and dimeric beta-bungarotoxin (β-BTx, 20 kDa), which form lighter monomers under reducing condition (Figure 2A). The chromatogram showed 17 major peaks, with peaks 8, 10 and 14 exhibiting the highest abundance (Figure 2B).

**Figure 2.**
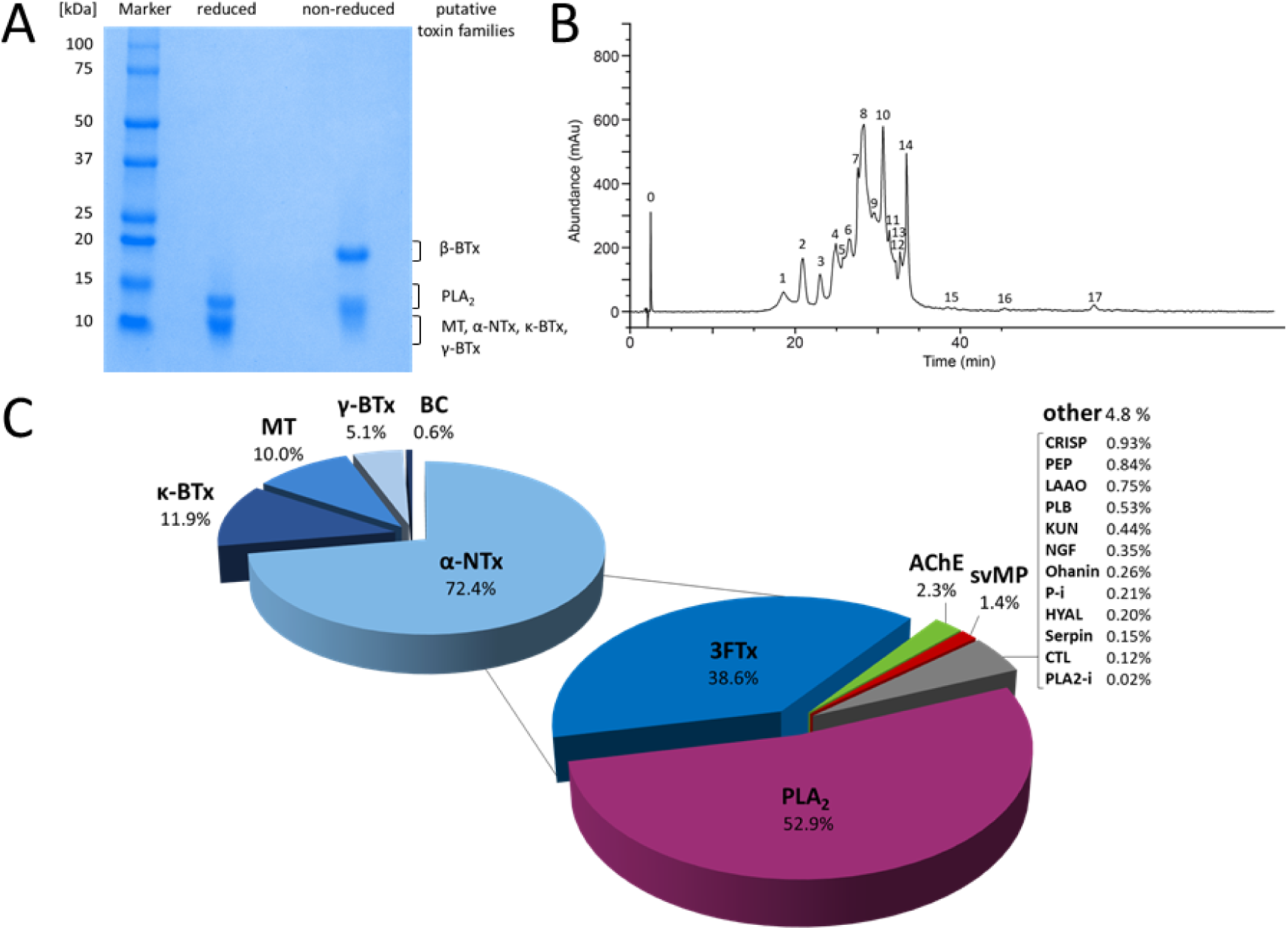
Characterization of *Bungarus multicinctus* venom. (A) One-dimensional SDS-PAGE profiles, (B) RP-HPLC profile and (C) *Bungarus multicinctus* venom composition based on bottom-up proteomics analysis. The relative abundances presented in the pie chart were calculated based on NSAFs. Abbreviations: 3FTx, three-finger-toxin; α-NTx, alpha-neurotoxin; κ-BTx, kappa-bungarotoxin; γ-BTx gamma-bungarotoxin; AChE, acetylcholinesterase; BC, bucandin; CRISP, cysteine-rich secretory protein; CTL, snake venom C-type lectins and C-type lectin-related proteins; HYAL, hyaluronidase; KUN, Kunitz-type serine protease inhibitors; LAAO, L-amino-acid-oxidase; P-i, protease-inhibitor; PLA_2_, phospholipase A_2_; PLA_2_-i, phospholipase A_2_-inhibitor; PLB, phospholipase B; svMP, snake venom metalloproteinase.

To provide a comprehensive atlas of the components within *B. multicinctus* venom we employed a previously sequenced high-quality genome produced for this species [40] as a peptide search database for bottom-up proteomics. Through this approach, we identified a total of 113 proteins, along with their nucleotide and amino acid sequences. Rigorous quality filtering, including BLAST searches, Interproscan, and manual comparative alignment investigation, allowed us to pinpoint 55 *bona fide* venom components categorized into 16 distinct protein families. These venom components and their primary structures are reported in supplementary Table S9. The most diverse protein family was 3FTx and peptidase with 14 entries each. Manual sequence analysis revealed, that the identified 3FTxs belong to five 3FTx-sub-classes: α-NTx, κ-BTx, γ-BTx, bucandin (BC) and MT.

To obtain a quantitative, and thus more realistic perspective on the *B. multicinctus* venom profile, we determined the relative abundance of each component by calculating their NSAF, which represents a reliable estimate of their relative abundance (Figure 2C). Our analysis revealed that PLA_2_ and 3FTx present the highest abundances (52.9% and 38.6%, respectively), accounting for over 90% of the *B. multicinctus* venom proteome. The third and fourth most abundant protein families were acetylcholinesterase (AChE; 2.3%) and snake venom metalloproteinase (svMP, 1.4%). Twelve additional protein families accounted in total for 4.8% of the venom proteome: cysteine-rich secretory proteins (CRISP; 0.9%), peptidase (PEP; 0.8%), L-amino-acid-oxidase (LAAO; 0.8%), phospholipase B-like-family (PLB; 0.5%), Kunitz-type serine-protease inhibitors (KUN; 0.4%), venom nerve growth factor (NGF; 0.4%), ohanin (0.3%), protease-inhibitor (P-i; 0.2%), hyaluronidase (HYAL; 0.2%), serpin (0.2%), snake venom C-type lectins and C-type lectin-related proteins (CTL; 0.1%) and phospholipase A_2_ inhibitor (PLA_2_-i; 0.02%). Additional information on the components identified and their abundances is reported in Table 1.

**Table 1:** Dimer composition of β-BTx in *B. multicinctus* venom. The table shows the observed and theoretical molecular weight, the mass error, as well as the putative chains of β-BTx from *B. multicinctus* venom as determined in top-down proteomic analysis.

| Obs. β-BTx [Da] | Theo. β-BTx [Da] | Mass error [ppm] | A chain (ID) | B chain (ID) |
| --- | --- | --- | --- | --- |
| 20648.38 | 20648.35 | 1.4 | A7 (Q9PU97) | B3 (Q9W728) |
| 20608.40 | 20608.34 | 2.8 | A3 (P00619) | B2 (P00989) |
| 20598.35 | 20598.15 | 9.3 | A4 (P17934) | B2 (P00989) |
| 20598.35 | 20599.30 | 46.2 | A-AL3 (Q9PTA6) | B5 (Q1RPS9) |
| 20598.35 | 20599.32 | 47.3 | A3 (P00619) | B1 (P00987) |
| 20598.35 | 20597.20 | 55.6 | A-AL3 (Q9PTA6) | B6 (Q1RPS8) |
| 20597.34 | 20597.20 | 6.8 | A-AL3 (Q9PTA6) | B6 (Q1RPS8) |
| 20597.34 | 20598.15 | 39.5 | A4 (P17934) | B3 (Q9W728) |
| 20591.36 | 20590.36 | 48.5 | A1 (P00617) | B4 (Q1RPT0) |

### 4.2 Verification of intact components and dimerization of β-bungarotoxins

To consider smaller venom components that are usually difficult to detect via traditional bottom-up proteomics, we supplemented our proteomic investigation by a top-down approach under non reduced and reduced condition (Figure S2).

This confirmed the presence of the previously determined major toxin families and enabled the identification of full-length sequences of 3FTx, KUN and PLA_2_, including the exact number of formed disulfide bridges (Table S13). In combination with the intact mass profiling, our analysis revealed that the non-reduced 20 kDa signals correspond to putative β-BTx that dissociate under reducing conditions into their constituent monomeric chains: a 13 kDa β-BTx A chain (PLA_2_) and a 7 kDa B chain (KUN). For example, the abundant 20608.40 Da β-BTx was shown (mass error of 2.8 ppm) to be formed by dimerization of the acidic PLA_2_ A3 chain (P00619, red. obs. 13415.93 Da) and the KUN homolog B2 chain (P00989, red. obs. 7192.41 Da). While the composition of some β-BTx isoforms was precisely determined, the high variability and close molecular masses of others indicated the presence of multiple possible combinations. (Table 1).

### 4.3 Bioactivity

After gaining a detailed insight into the components of *B. multicinctus* venom, we aimed to assess the functional relevance of the obtained compositional data. To this end, we performed a broad array of bioassays targeting a range of biological activities possibly exerted by *B. multicintus* venom.

To investigate the biological activity of *B. multicinctus* venom, we conducted PLA_2_, AChE, and protease activity assays. In all performed assays we observe enzymatic activity in a positively correlated concentration-dependent manner. *Bungarus multicinctus* venom showed high levels of PLA_2_ activity across all measured concentrations. At 3.125 µg/ml the PLA_2_ activity was 41.1%, and 70.6% at 6.25 µg/ml. Starting at concentration 12.5 µg/ml the venom surpassed the positive control, therefore exceeding 100% (Figure 3A). The AChE activity was very high in all measured concentrations exceeding the calibrator (equivalent to 200 units/l) even in the lowest concentration. The highest activity was observed at 5 µg/ml with 927.0 Units/l, followed by 2.5 µg/ml with 505.0 Units/l, and the lowest at 1.25 µg/ml with 261.2 Units/l (Figure 3B). The venom showed only low levels of protease activity across all tested concentrations, ranging between 3.0% (50 µg/ml) and 5.2% (200 µg/ml) (Figure 3C).

**Figure 3.**
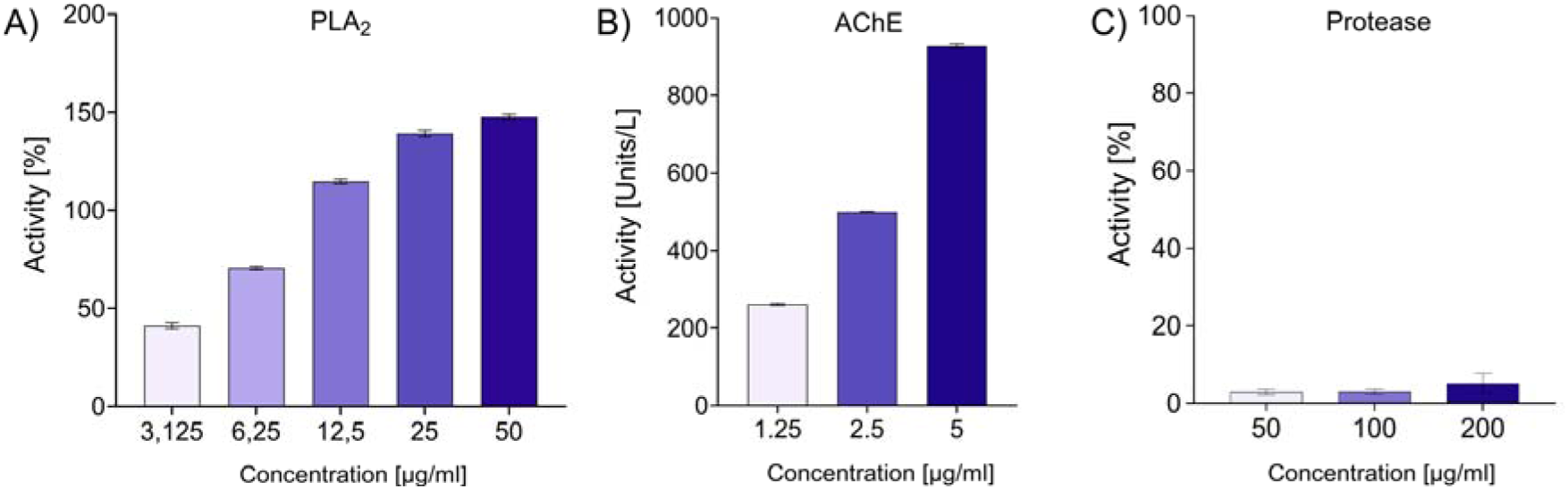
Enzymatic activity of *B. multicinctus* venom. (A) Phospholipase A_2_ (PLA_2_), (B) Acetylcholinesterase (AChE) and (C) Protease activities were tested at different venom concentrations. The values presented in each graph refer to the normalized averages of the performed measurements.

To further investigate the clinical relevance of the analyzed venom we conducted cell viability assays. Specifically, we tested the following cell lines: PMBC, HEK293T, RAW 264.7, SH-SY5Y at various concentrations. None of the tested venom concentrations showed cytotoxic effects against any of the cell lines considered at any tested concentration (Figure 4).

**Figure 4.**
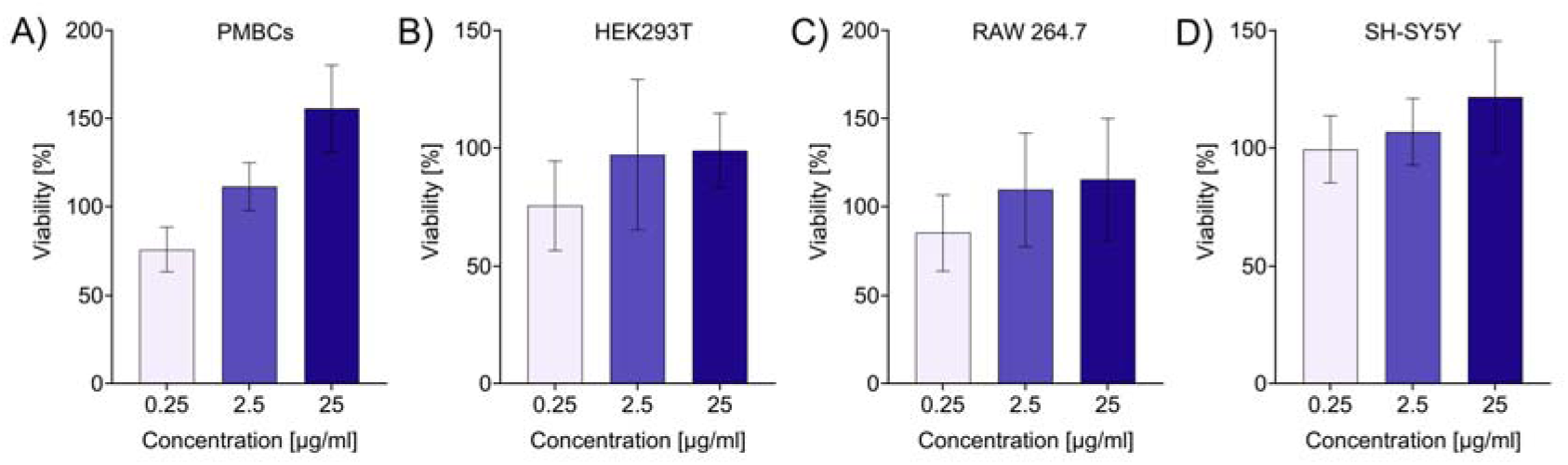
Cytotoxic activity of *B. multicinctus* venom. The graphs show the normalized values of cell viability of the different cell lines tested (PMBCs, HEK293T, RAW 264.7, SH-SY5Y) at three different venom concentrations (0.25, 2.5, 25 µg/ml). Error mean in ±SD.

Following our investigation of enzymatic activities and in vitro assays on cell cultures, we proceeded to study additional, less commonly examined effects from snake venom. As krait venom is known to often exert antimicrobial and antiviral activity [64], we opted to explore these activities in our sample. First, we screened *B. multicinctus* venom against a variety of clinically or environmentally important bacteria. Only minor antibacterial effects were observed on the Gram-negative bacteria tested, with the strongest antibacterial effect detected against *E. coli* at the higher venom concentrations (2, 1, 0.5 mg/ml), reducing the bacterial growth rate to ∼33%. Stronger effects were exhibited against the tested Gram-positive bacteria, excluding *S. epidermidis,* where no antibacterial effect could be observed. The strongest antibacterial effect was seen in *S. aureus*, where the bacteria growth was reduced to ∼0% at the three higher venom concentrations (2, 1, 0.5 mg/ml), despite no antibacterial effect at the two lower concentrations (0.25, 0.125 mg/ml). A similar pattern was detected in *M. luteus*, where only in the higher concentrations (2, 1, 0.5 mg/ml) an antibacterial effect was seen. In contrast an antibacterial effect against *L. monocytogenes* could be observed in a postive concentration-dependent manner, with growth rates ranging from 32 to 79% (Figure 5). Furthermore, we conducted an antiviral activity assay of *B. multicinctus* against three clinically important influenza virus strains. The venom showed no antiviral activity at any tested concentration (Figure S1).

**Figure 5.**
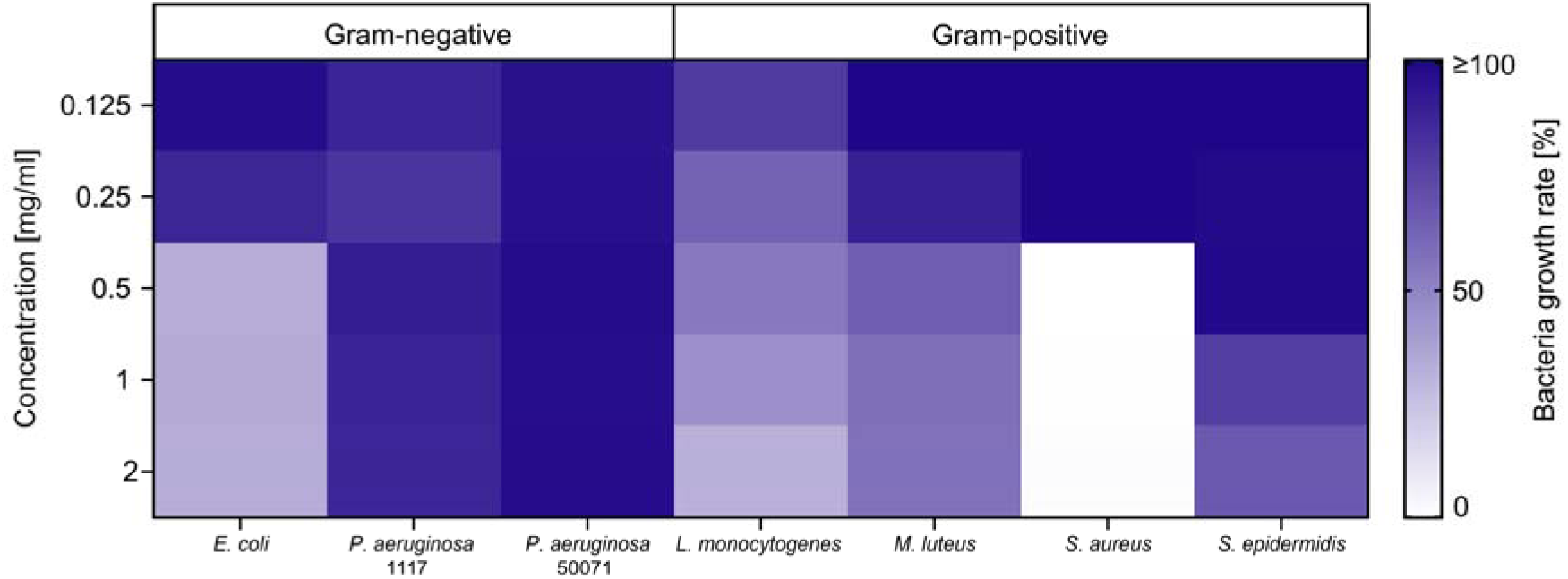
Antibacterial activity of *B. multicinctus* venom. The heatmap shows the photometrically determined bacterial growth rate [%] of three Gram-negative and four Gram-positive strains after exposure to different venom concentrations. The values presented refer to the normalized averages of the values of three measurements per concentration.

Finally, we conducted assays to explore effects on the release of second messengers following exposure to *B. multicinctus* venoms. We carried out NO assays utilizing RAW264.7 cells investigating the NO-induction and inhibition after 24 h, as well as Ca^2+^-assays using HEK293T cells where we looked into Ca^2+^-release inhibition and induction. Overall, the analyzed *B. multicinctus* venom showed no effect on intracellular Ca^2+^ release and NO levels (Table S5, S6)

## 5 Discussion

### 5.1 The first genome-annotated venom proteome of *Bungarus multicinctus*

The increasing size, availability, and quality of next-generation databases and the resulting growing extent in their utilization, offers solutions to various important obstacles of proteomic inferences [65]. Through extensive proteogenomics utilizing a previously sequenced high-quality genome produced for *B. multicinctus* [40] in combination with protein profiling as well as bottom-up and top-down proteomics we were able to present the first genome-based atlas of the *B. multicinctus* venom arsenal.

This analysis revealed an overall quite simple venom that contains a total 55 *bona fide* toxin proteins, categorized into 16 groups, for which we successfully retrieved the primary structures from the genome with highest confidence (Table S9, S11). The two toxin families 3FTx and PLA_2_ make up a total of 91.5% of the quantitative *B. multicinctus* venom proteome, aligning with the compositional pattern commonly observed in most elapid venoms, particularly within the genus *Bungarus* [11,13,66,67].

When comparing our work to other proteomes of *B. multicinctus*, we detect evident similarities as well as some interesting differences to previous works. For instance, the studies by Shan *et al.* 2016 and Oh *et al*. 2021 identified the same major components, although with different abundances, particularly concerning 3FTx and PLA_2_. While in both previous works the venom mostly contained 3FTx (∼60-77%), our results showed a higher abundance of PLA_2_ (52.9%), while 3FTx featured a smaller fraction of the venom proteome (38.6%) (Figure 2C)[37,68]. Furthermore, we noticed that many minor components (e.g. PEP, PLB and KUN) were not reported. Additional discrepancies can be observed considering the number of components detected across studies. For instance, Oh et al. (2021) identified only 36 proteins from seven distinct toxin families in their *B. multicinctus* venom proteome from Taiwan [37]. Likewise, Shan et al. (2016) reported only seven protein families (3FTx, PLA_2_, KUN, svMP, LAAO, AChE and PEP) in the analyzed Chinese *B. multicinctus* venoms [68]. Lastly, another *B. multicinctus* venom proteome sourced from Vietnamese specimen reported 17 protein families [35]. That said, while our generated *B. multicinctus* venom proteome contains a much higher diversity of venom components than most of these previous works, it must be noted that the major venom components identified are in agreement across the so far conducted surveys. All works reported a venom dominated by 3FTx and PLA_2_, and also containing KUN, svMP, LAAO, AChE and NGF in noteworthy amounts. They further list selected minor protein groups (e.g. PEP, PLB, ohanin).

The previously generated proteomes for different *B. multicinctus* venoms were published at different time points across the last decade and by researchers from different laboratories. Hence, they likely relied on different mass spectrometric platforms, quantification approaches and distinct databases used for protein identification. In this context, the observed discrepancies are at least partly attributable to the application of different methodological approaches, hindering for instance the possibility for direct quantitative comparisons. Yet, the consensus of all thus far analyzed *B. multicinctus* venoms suggest a largely PLA_2_ and 3FTx dominated venom, although their ratios may differ between venom samples. Such variations were already discussed previously and could be due to intraspecific venom variation [35,37,68].

In general venom variation in *Bungarus* has received comparatively little attention thus far. However, a recent functional venomics analysis of the Indian krait (*Bungarus caeruleus*) across India revealed tremendous differences in venom compositions and activities between geographically distinct populations of that species [69]. These are reminiscent to the patterns observed when comparing *B. multicinctus* proteomes from different studies (often reflecting different localities). The work on *B. caeruleus* revealed strong differences in nAChR binding properties and alarming discrepancies in their ability to be neutralized by commercially available antivenoms, some of which virtually fail against certain populations [69]. Most interesting in this study is the apparent presence of venom variation also on the proteomic level, with PLA_2_ ranging from 45% - 67%, 3FTx ranging from 10-29%, and β-BTx representing 6-14% between investigated populations [69]. In light of the pioneering insights from *B. caeruleus*, we believe that the herein revealed variability in *B. multicinctus* venom profiles suggests a noteworthy degree of geographic venom variation in this species and potentially within kraits in general. Therefore, future studies should aim at increasing our knowledge on the variability on *Bungarus* venoms, in *B. multicinctus* and beyond, by investigating a larger sample size of localities and hitherto unstudied species. However, it has also been shown that cryptic speciation also contributes to aberrant venom profiles in kraits and that overlooked species boundaries may blur our understanding of this phenomenon [70]. This could also affect *B. multicinctus*, which features a species with unclear delimitation and potentially containing cryptic species, as proposed via the *Bungarus candidus/multicinctus/wanghaotingi* complex where lineages are genetically and morphologically difficult to distinguish from closely related species with similar black-white-crossbands [71]. Here, demarcating the boundaries of *B. multicinctus* and *B. wanghaotingi* is particularly challenging because of their strong similarities [71,72]. Multiple distinct but yet undescribed species are actually suspected to belong to the “*multicinctus-wanghaotingi*” clade, probably being polytypic with several distinct species being hidden under this clade name, hence deserving more thorough investigation [73].

### 5.2 Complex dimerization patterns of β-bungarotoxins

Neurotoxicity is the major contributor to the high mortality rate following envenomation by kraits [74–76]. Among the main active principles behind the neurotoxic symptoms elicited are β-bungarotoxins. These heterodimeric toxins, consisting of PLA_2_ and KUN, target presynaptic motor nerve terminals and destroy synaptic vesicles, causing permanent damage to the nerves [74,75,77]. This results in long-lasting flaccid paralysis of skeletal muscles, leaving krait snakebite victims on ventilation until the nerve terminals are regenerated [29,38,74]. Given the medical relevance of bungarotoxins, several studies have extensively investigated their structure and occurrence across the genus *Bungarus*. However, remarkably little information is available on β-BTx. Particularly, it is not known if the pairing of different proteoforms may lead to activity changes, and whether each subunit chain can freely dimerize with each other has yet to be investigated.

By comparing the non-reduced and reduced SDS-PAGE and top-down profiles of *B. multicinctus* venom, we were able to confirm the presence of β-BTx with different A and B chain combinations. Our results showed five A- and five B chains in ten different combinations, reflecting only the most abundant β-BTx signals in the analyzed venom, providing a first insight on how variable A and B chain combinations could be. There are only few other reports of A and B chain combinations in β-BTx, indicating that while some A chain PLA_2_s and B chain KUNs have often been identified within β-BTx, others are detected less frequently [78]. Beside β-BTx, the members of several other snake toxin families are known to multimerize, including multimeric CTL, dimeric disintegrins, as well as P-III svMP-CTL complexes [79]. To date, the effects induced by subunit-alteration in most of these toxins are still unknown and may represent promising avenues for the development of novel therapeutic tools for the management and treatment of snakebite.

### 5.3 Novel insights into the pathophysiology of venom-induced neurotoxicity

The pathophysiological effects caused by *B. multicinctus* envenoming consist mainly of ptosis, dysarthria, dilated pupils and peripheral paralysis, which can ultimately progress to respiratory paralysis and death within 1.5–6.5 hours [28,29,38,80]. The occurrence of such strong neurotoxic effects, often accompanied by general pain but without local symptoms at the bite site [4,29,38,81], is consistent with the high abundances of neurotoxins present in this species’ venom, such as 3FTx and PLA_2_ [35,37,68](Figure 2C).

Given the medical significance of snake venom PLA_2_, as well as the high abundance of this toxin family in our venom proteome, we aimed to investigate the PLA_2_ activity of the analyzed *B. multicintus* venom. The performed PLA_2_ activity bioassay showed high potency of the venom across all tested concentrations, even surpassing the positive control at the three highest concentrations (Figure 3A). Considering that PLA_2_s, particularly beta-bungarotoxin, are the main cause of flaccid paralysis and long-lasting neurological symptoms after *B. multicinctus* envenomation [74–76], the high PLA_2_ activity recorded is consistent with the strong neurotoxicity typically reported after many-banded krait bites [29,38]. Nonetheless, while potent PLA_2_ activity was evident *in vitro*, further tests (ideally *in vivo*) are required to confirm its role in the neurotoxic effects of *B. multicinctus* snakebite.

Another venom component potentially involved in the neurotoxic effects following many-banded krait envenomation is AChE. While there is still uncertainty concerning the actual toxicity of AChE in terms of snakebite, both depletion and accumulation of this enzyme at the neuromuscular junction can be harmful, potentially leading to either flaccid or tetanic paralysis [82]. Furthermore, AChE is commonly found in many elapid venoms, and the venoms of several *Bungarus* species are known to exhibit high AChE activity [82], potentially indicating a role in envenomation. In light of this, and considering that the relative abundance of AChE in the analyzed venom (2.3%) is markedly higher than that reported for other *B. multicinctus* venom proteomes (ranging from 0.02% to 0.5%) [35,37,68], we measured the AChE activity of our venom sample in vitro. Similarly, to what we observed for PLA_2_, the analyzed *B. multicinctus* venom presented remarkably high AChE activity, even at low concentrations (Figure 3B). While these findings suggest that AChE may contribute to the envenomation profile of *B. multicinctus*, in vivo studies involving living models (e.g., natural prey items) would be needed to elucidate the functional role of this potent activity.

In line with the absence of local symptoms in *Bungarus* snakebite victims, many-banded krait venoms typically contain little to no proteases (e.g., svMP, svSP), components commonly associated with local effects like swelling and necrosis at the bite site [29,38,81,83,84]. Nonetheless, as svMPs were detected in the venom proteome (∼1.4% of the total composition), we aimed to assess their potency and relevance to envenomation by evaluating the protease activity and cytotoxicity of *B. multicinctus* venom. The performed bioassays retrieved virtually no proteolytic activity, as well as no cytotoxic effects on any of the four tested cell lines at any concentration (Figure 3C; 4). While these results are unsurprising given the lack of clinical manifestations attributable to proteolytic mechanisms or cytotoxicity in many-banded krait envenomation, they allow us to foster our understanding on the pathophysiology of its neurotoxicity. Several components, including β-bungarotoxins, are well known to exert their neurotoxic effects through damage at the nerves [77,85]. Hence, we specifically investigated cell viability of SH-SY5Y, a neuronal cell line often used to study neuronal function and differentiation, after exposure to *B. multicinctus* venom. Similar to other tested cells, no effect on cell viability was detected at any concentration despite the damage that some *B. multicinctus* venom components cause at nerve cells. Therefore, it appears that neurons are actively targeted by the venom and functionally impaired on several levels, some of which causing damage to the cell. However, these effects do not lead to the death of neuronal cells. This could explain why patients bitten by kraits are able to recover after antivenom treatment and receiving ventilation, although the recovery can be a slow process [29,38]. Further, our analysis of second messengers revealed, that *B. multicinctus* venom does not interfere with their release inhibition or induction. Hence, at least these pathways are supposedly not involved in the manifestation of many-banded krait neurotoxicity on the cellular level.

## 6 Conclusion and future perspectives

The many-banded krait, *B. multicinctus*, represents one of Earth’s most lethal and infamous snakes. Yet, surprisingly little is known about its venom in comparison to other snakes of similar medical and cultural impact.

Here, we provide the first proteogenomic and functional assessment of its venom and report its complete compositional profile, the primary structures of its constituents, the complex dimerization landscape of β-bungarotoxins, and its activity profile. We show that the venom is largely composed by 3FTx and PLA_2_ and causes highly potent PLA_2_ as well as AChE activities, suggesting major roles of these compounds in developing neurotoxic symptoms. We recommend that a detailed characterization of these enzymes should be carried out to better understand their role in the pathophysiology of *B. multicinctus* venoms. The conduction of in vitro assays on various cell lines further added to our understanding of *B. multicinctus* envenoming. The absence of cytotoxicity was unsurprising when taking the clinical effects of bites into account. Yet, we also detected no effect on the viability of neuronal cells suggesting that, despite the damage caused on neurons by several krait toxins, their viability is not impaired [74,75,77]. We further found no evidence that *B. multicinctus* venom interferes with second messenger release. However, both of these findings require further validation, optimally using in vivo systems. Interestingly, potential medical applications are often proposed for krait venoms, particularly in the areas of antiinfectives and anticancer agents [64]. In our functional assessment, we therefore tested the effects of *B. multicinctus* venom on various bacterial and viral (influenza) strains. While we retrieved no activity against the tested viruses, we found potent growth inhibitory effects against *Staphylococcus aureus*. Thus, it might be promising to explore *B. multicinctus* venom against a larger diversity of clinically relevant and drug-resistant strains of *S. aureus* to identify potential antibiotic leads.

Lastly, when comparing our venom proteome with previously published datasets, we noticed some interesting differences. While virtually all *B. multicinctus* venom proteomes agree on the same major components (essentially 3FTx and PLA_2_), their ratios appear to differ between studies. While this might, at least partially, be attributable to methodological bias, our findings may likewise pinpoint to the presence of intraspecific venom variation. This phenomenon occurs widely in snakes and has only recently been reported in kraits [69,70]. In order to resolve this question, it will be necessary to carry out a strategic survey of venoms from *B. multicinctus* venoms from ecologically and phylogenetically distinct populations from across its distribution range and test them under unified experimental conditions. As intraspecific venom variation could also stem from the presence of cryptic species [70], it is important to also investigate and appropriately consider species boundaries within examined *Bungarus* (*multicinctus*) venoms. Further, it will be important to investigate the species natural history in more detail in order to establish a framework in which the potentially occurring venom variation can be interpreted in a meaningful way.

Overall, we consider the available data on *B. multicinctus* venom proteomes and its congeners justifies the assumption that intraspecific venom variation is more common in kraits than currently acknowledged. We believe it is therefore of pivotal importance to investigate this phenomenon in this medically important genus with velocity as is currently done in other highly medically important snakes (e.g. the genera *Echis*, *Daboia* and *Naja*).

## Acknowledgments

The authors acknowledge various members of the German Society for Herpetology and Herpetoculture for their insightful discussions on krait biology that influenced this manuscript. We further thank Benno Kreuels (Bernhard Nocht Institute for Tropical Medicine) for introducing several participants of this study to each other and therefore for creating the fundamental base for this work.

## Author contributions (CRediT)

**Lilien Uhrig**: Writing – review & editing, Writing – original draft, Visualization, Supervision, Project administration, Methodology, Investigation, Formal analysis, Data curation, Conceptualization. **Ignazio Avella**: Writing – review & editing, Visualization, Software, Methodology, Investigation, Formal analysis, Data curation. **Lennart Schulte**: Writing – review & editing, Visualization, Software, Methodology, Investigation, Formal analysis, Data curation. **Sabine Hurka**: Writing – review & editing, Visualization, Software, Methodology, Investigation, Formal analysis, Data curation. **Ludwig Dersch**: Writing – review & editing, Investigation, Formal analysis, Data curation. **Johanna Eichberg**: Writing – review & editing, Investigation, Formal analysis, Data curation. **Susanne Schiffmann**: Writing – review & editing, Investigation, Formal analysis, Data curation. **Marina Henke**: Investigation, Formal analysis, Data curation. **Ulrich Kuch**: Writing – review & editing, Resources. **Alfredo Cabrera-Orefice**: Writing – review & editing, Software, Resources, Methodology, Investigation, Formal analysis, Data curation. **Kornelia Hardes**: Writing – review & editing, Resources. **Andreas Vilcinskas**: Writing – review & editing, Supervision, Project administration, Funding acquisition. **Maik Damm**: Writing – review & editing, Visualization, Software, Methodology, Supervision, Formal analysis, Investigation, Data curation. **Tim Lüddecke**: Writing – review & editing, Visualization, Supervision, Software, Resources, Project administration, Methodology, Investigation, Funding acquisition, Formal analysis, Data curation, Conceptualization

## 8 Supplementary Information

**Figure S1.**
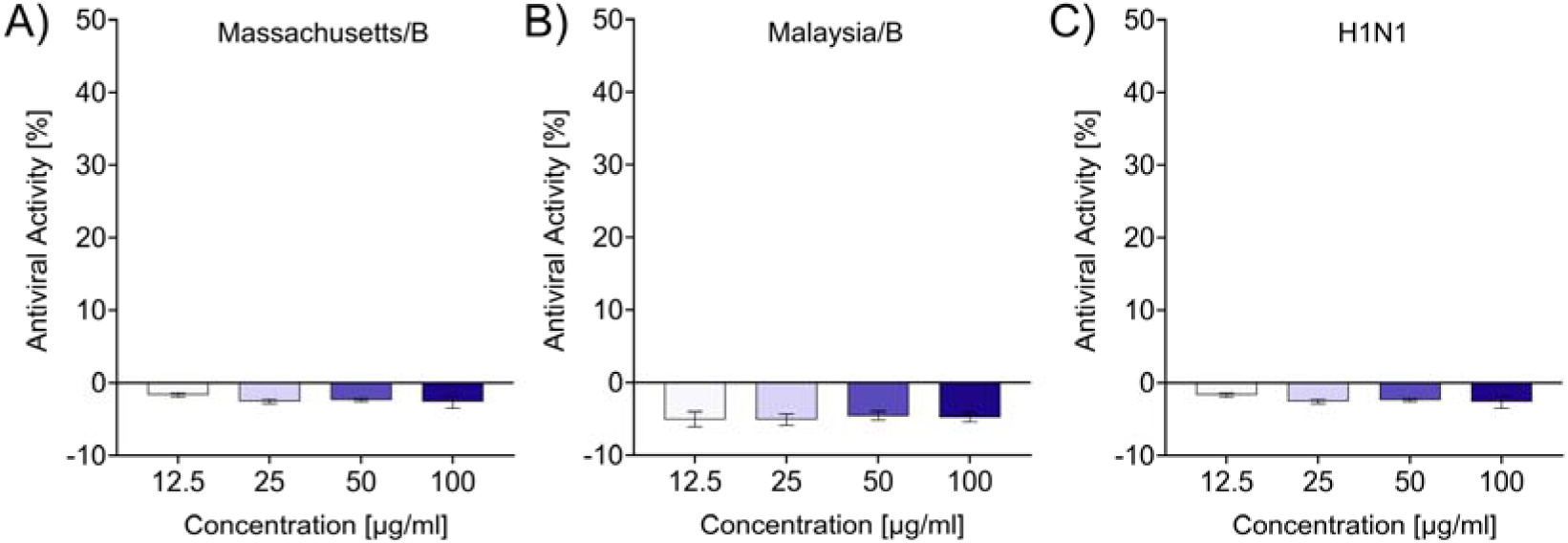
Antiviral activity of *B. multicinctus* venom against virus-infected MDCK II cells. Three different influenza strains (Massachusetts/B (MOI□=□0.01), B/Malaysia/2506/2004 (Malaysia/B; MOI=1) and A/Hamburg/05/2009 (H1N1; MOI=1)) used to infect canine kidney cells (MDCK II). Raw luminescence values were baseline-corrected to the virus control (0%) and normalized to the aprotinin-positive control (100%). Data are means□±□SD of n =□3 independent measurements.

**Figure S2.**
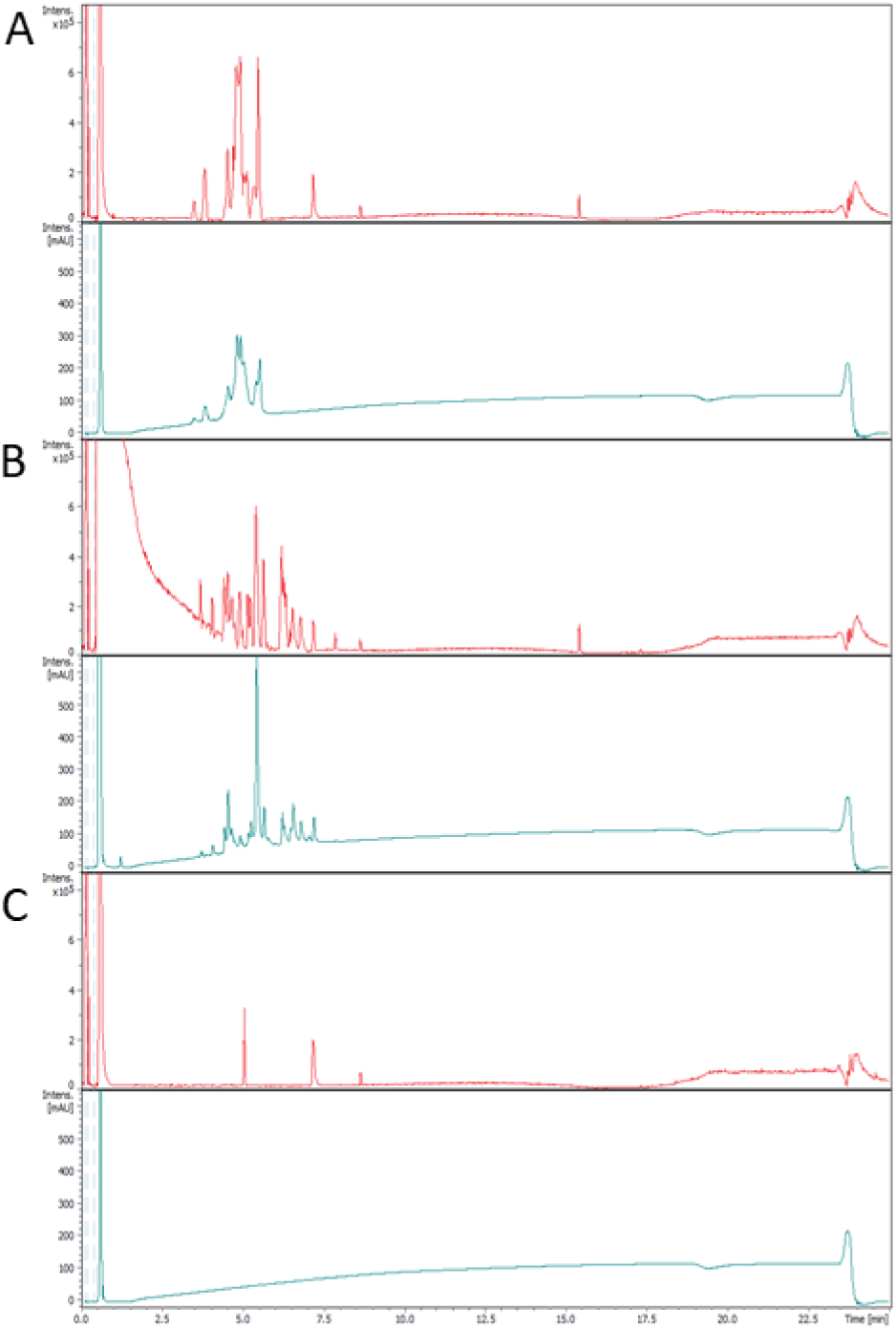
Top-down venom profile of *Bungarus multicinctus*. For top-down and intact mass profiling the venom profile was detected under (A) non-reducing and (B) TCEP reducing conditions (with a strong TCEP signal at early retention times) as well as (C) the pre-analysis blank. With (red, top) base peak current (BOC) and simultaneously the (blue, bottom) RP-HPLC at λ = 214 nm for each sample.

## Bibliography

[1] R.A. Harrison, A. Hargreaves, S.C. Wagstaff, B. Faragher, D.G. Lalloo, Snake Envenoming: A Disease of Poverty, PLoS Negl. Trop. Dis. 3 (2009) e569. 10.1371/journal.pntd.0000569.

[2] J.-P. Chippaux, Estimate of the burden of snakebites in sub-Saharan Africa: a meta-analytic approach, Toxicon Off. J. Int. Soc. Toxinology 57 (2011) 586–599. 10.1016/j.toxicon.2010.12.022.

[3] B. Mohapatra, D.A. Warrell, W. Suraweera, P. Bhatia, N. Dhingra, R.M. Jotkar, P.S. Rodriguez, K. Mishra, R. Whitaker, P. Jha, Million Death Study Collaborators, Snakebite mortality in India: a nationally representative mortality survey, PLoS Negl. Trop. Dis. 5 (2011) e1018. 10.1371/journal.pntd.0001018.

[4] J.M. Gutiérrez, J.J. Calvete, A.G. Habib, R.A. Harrison, D.J. Williams, D.A. Warrell, Snakebite envenoming, Nat. Rev. Dis. Primer 3 (2017) 1–21. 10.1038/nrdp.2017.63.

[5] R.W. Snow, R. Bronzan, T. Roques, C. Nyamawi, S. Murphy, K. Marsh, The prevalence and morbidity of snake bite and treatment-seeking behaviour among a rural Kenyan population, Ann. Trop. Med. Parasitol. 88 (1994) 665–671. 10.1080/00034983.1994.11812919.

[6] A. Kasturiratne, A.R. Wickremasinghe, N. de Silva, N.K. Gunawardena, A. Pathmeswaran, R. Premaratna, L. Savioli, D.G. Lalloo, H.J. de Silva, The Global Burden of Snakebite: A Literature Analysis and Modelling Based on Regional Estimates of Envenoming and Deaths, PLOS Med. 5 (2008) e218. 10.1371/journal.pmed.0050218.

[7] World Health Organization, Snakebite envenoming: a strategy for prevention and control, World Health Organization, Geneva, 2019. https://iris.who.int/handle/10665/324838 (accessed April 16, 2025).

[8] D.J. Williams, M.A. Faiz, B. Abela-Ridder, S. Ainsworth, T.C. Bulfone, A.D. Nickerson, A.G. Habib, T. Junghanss, H.W. Fan, M. Turner, R.A. Harrison, D.A. Warrell, Strategy for a globally coordinated response to a priority neglected tropical disease: Snakebite envenoming, PLoS Negl. Trop. Dis. 13 (2019) e0007059. 10.1371/journal.pntd.0007059.

[9] B.M. von Reumont, G. Anderluh, A. Antunes, N. Ayvazyan, D. Beis, F. Caliskan, A. Crnković, M. Damm, S. Dutertre, L. Ellgaard, G. Gajski, H. German, B. Halassy, B.-F. Hempel, T. Hucho, N. Igci, M.P. Ikonomopoulou, I. Karbat, M.I. Klapa, I. Koludarov, J. Kool, T. Lüddecke, R. Ben Mansour, M. Vittoria Modica, Y. Moran, A. Nalbantsoy, M.E.P. Ibáñez, A. Panagiotopoulos, E. Reuveny, J.S. Céspedes, A. Sombke, J.M. Surm, E.A.B. Undheim, A. Verdes, G. Zancolli, Modern venomics—Current insights, novel methods, and future perspectives in biological and applied animal venom research, GigaScience 11 (2022) giac048. 10.1093/gigascience/giac048.

[10] T. Lüddecke, A. Paas, R.J. Harris, L. Talmann, K.N. Kirchhoff, A. Billion, K. Hardes, A. Steinbrink, D. Gerlach, B.G. Fry, A. Vilcinskas, Venom biotechnology: casting light on nature’s deadliest weapons using synthetic biology, Front. Bioeng. Biotechnol. 11 (2023). 10.3389/fbioe.2023.1166601.

[11] A.L. Oliveira, M.F. Viegas, S.L. da Silva, A.M. Soares, M.J. Ramos, P.A. Fernandes, The chemistry of snake venom and its medicinal potential, Nat. Rev. Chem. 6 (2022) 451–469. 10.1038/s41570-022-00393-7.

[12] B.G. Fry, N.R. Casewell, W. Wüster, N. Vidal, B. Young, T.N.W. Jackson, The structural and functional diversification of the Toxicofera reptile venom system, Toxicon 60 (2012) 434–448. 10.1016/j.toxicon.2012.02.013.

[13] T. Tasoulis, G.K. Isbister, A Review and Database of Snake Venom Proteomes, Toxins 9 (2017) 290. 10.3390/toxins9090290.

[14] G. Zancolli, J.J. Calvete, M.D. Cardwell, H.W. Greene, W.K. Hayes, M.J. Hegarty, H.-W. Herrmann, A.T. Holycross, D.I. Lannutti, J.F. Mulley, L. Sanz, Z.D. Travis, J.R. Whorley, C.E. Wüster, W. Wüster, When one phenotype is not enough: divergent evolutionary trajectories govern venom variation in a widespread rattlesnake species, Proc. R. Soc. B Biol. Sci. 286 (2019) 20182735. 10.1098/rspb.2018.2735.

[15] N.R. Casewell, T.N. Jackson, A.H. Laustsen, K. Sunaga, Causes and Consequences of Snake Venom Variation, Trends Pharmacol. Sci. 41 (2020) 570. 10.1016/j.tips.2020.05.006.

[16] M. Damm, B.-F. Hempel, R.D. Süssmuth, Old World Vipers—A Review about Snake Venom Proteomics of Viperinae and Their Variations, Toxins 13 (2021) 427. 10.3390/toxins13060427.

[17] I. Avella, W. Wüster, L. Luiselli, F. Martínez-Freiría, Toxic Habits: An Analysis of General Trends and Biases in Snake Venom Research, Toxins 14 (2022) 884. 10.3390/toxins14120884.

[18] W. Rao, K. Kalogeropoulos, M.E. Allentoft, S. Gopalakrishnan, W. Zhao, C.T. Workman, C. Knudsen, B. Jiménez-Mena, L. Seneci, M. Mousavi-Derazmahalleh, T.P. Jenkins, E. Rivera-de-Torre, S. Liu, A.H. Laustsen, The rise of genomics in snake venom research: recent advances and future perspectives, GigaScience 11 (2022) giac024. 10.1093/gigascience/giac024.

[19] K. Sunagar, D. Morgenstern, A.M. Reitzel, Y. Moran, Ecological venomics: How genomics, transcriptomics and proteomics can shed new light on the ecology and evolution of venom, J. Proteomics 135 (2016) 62–72. 10.1016/j.jprot.2015.09.015.

[20] S.H. Drukewitz, B.M. von Reumont, The Significance of Comparative Genomics in Modern Evolutionary Venomics, Front. Ecol. Evol. 7 (2019). 10.3389/fevo.2019.00163.

[21] S.L. Thornton, Snakes, in: P. Wexler (Ed.), Encycl. Toxicol. Third Ed., Academic Press, Oxford, 2014: pp. 310–312. 10.1016/B978-0-12-386454-3.00786-7.

[22] Clinical Toxinology in Asia Pacific and Africa | SpringerLink, (n.d.). https://link.springer.com/referencework/10.1007/978-94-007-6386-9 (accessed April 21, 2025).

[23] Snakebite Information and Data Platform, (n.d.). https://www.who.int/teams/control-of-neglected-tropical-diseases/snakebite-envenoming/snakebite-information-and-data-platform (accessed January 29, 2025).

[24] Bungarus multicinctus, Reptile Database (n.d.). https://reptile-database.reptarium.cz/species.php?genus=Bungarus&species=multicinctus (accessed April 21, 2025).

[25] Search results | The Reptile Database, (n.d.). https://reptile-database.reptarium.cz/advanced_search?taxon=Elapidae&submit=Search (accessed November 13, 2024).

[26] P. Gopalkrishnakone, L.M. Chou, Snakes of Medical Importance: Asia-Pacific Region, Venom and Toxin Research Group, National University of Singapore, 1990.

[27] J.-J. Mao, G. Norval, C.-L. Hsu, W.-H. Chen, Ray Ger, Observations and comments on the diet of the many-banded krait (Bungarus multicinctus multicinctus) in Taiwan., Reptil. Amphib. Vol 17, No 2 (2010) 73–76.

[28] T. Pe, T. Myint, A. Htut, T. Htut, A.A. Myint, N.N. Aung, Envenoming by Chinese krait (Bungarus multicinctus) and banded krait (B. fasciatus) in Myanmar, Trans. R. Soc. Trop. Med. Hyg. 91 (1997) 686–688. 10.1016/S0035-9203(97)90524-1.

[29] Y.-C. Mao, P.-Y. Liu, L.-C. Chiang, S.-C. Liao, H.-Y. Su, S.-Y. Hsieh, C.-C. Yang, Bungarus multicinctus multicinctus Snakebite in Taiwan, Am. J. Trop. Med. Hyg. 96 (2017) 1497–1504. 10.4269/ajtmh.17-0005.

[30] C.A. Ariaratnam, M.H.R. Sheriff, R.D.G. Theakston, D.A. Warrell, Distinctive Epidemiologic and Clinical Features of Common Krait (Bungarus caeruleus) Bites in Sri Lanka, Am. J. Trop. Med. Hyg. 79 (2008) 458–462. 10.4269/ajtmh.2008.79.458.

[31] S.A.M. Kularatne, Common krait (Bungarus caeruleus) bite in Anuradhapura, Sri Lanka: a prospective clinical study, 1996–98, Postgrad. Med. J. 78 (2002) 276–280. 10.1136/pmj.78.919.276.

[32] R.K. Saini, S. Singh, S. Sharma, V. Rampal, A.S. Manhas, V.K. Gupta, Snake bite poisoning presenting as early morning neuroparalytic syndrome in jhuggi dwellers, J. Assoc. Physicians India 34 (1986) 415–417.

[33] W. Suraweera, D. Warrell, R. Whitaker, G. Menon, R. Rodrigues, S.H. Fu, R. Begum, P. Sati, K. Piyasena, M. Bhatia, P. Brown, P. Jha, Trends in snakebite deaths in India from 2000 to 2019 in a nationally representative mortality study, ELife 9 (2020) e54076. 10.7554/eLife.54076.

[34] D.P. Pandey, P. Bhattarai, R.C. Piya, Food Spectrum of Common Kraits (Bungarus caeruleus): An Implication for Snakebite Prevention and Snake Conservation, J. Herpetol. 54 (2020) 87–96. 10.1670/18-054.

[35] R.H. Ziganshin, S.I. Kovalchuk, G.P. Arapidi, V.G. Starkov, A.N. Hoang, T.T. Thi Nguyen, K.C. Nguyen, B.B. Shoibonov, V.I. Tsetlin, Y.N. Utkin, Quantitative proteomic analysis of Vietnamese krait venoms: Neurotoxins are the major components in *Bungarus multicinctus* and phospholipases A2 in *Bungarus fasciatus*, Toxicon 107 (2015) 197–209. 10.1016/j.toxicon.2015.08.026.

[36] L.-L. Shan, J.-F. Gao, Y.-X. Zhang, S.-S. Shen, Y. He, J. Wang, X.-M. Ma, X. Ji, Proteomic characterization and comparison of venoms from two elapid snakes (*Bungarus multicinctus* and *Naja atra*) from China, J. Proteomics 138 (2016) 83–94. 10.1016/j.jprot.2016.02.028.

[37] A.M.F. Oh, K.Y. Tan, N.H. Tan, C.H. Tan, Proteomics and neutralization of *Bungarus multicinctus* (Many-banded Krait) venom: Intra-specific comparisons between specimens from China and Taiwan, Comp. Biochem. Physiol. Part C Toxicol. Pharmacol. 247 (2021) 109063. 10.1016/j.cbpc.2021.109063.

[38] H.T. Hung, J. Höjer, N.T. Du, Clinical features of 60 consecutive ICU-treated patients envenomed by Bungarus multicinctus, Southeast Asian J. Trop. Med. Public Health 40 (2009) 518–524.

[39] Z.-Y. Zhang, Y. Lv, W. Wu, C. Yan, C.-Y. Tang, C. Peng, J.-T. Li, The structural and functional divergence of a neglected three-finger toxin subfamily in lethal elapids, Cell Rep. 40 (2022) 111079. 10.1016/j.celrep.2022.111079.

[40] J. Xu, S. Guo, X. Yin, M. Li, H. Su, X. Liao, Q. Li, L. Le, S. Chen, B. Liao, H. Hu, J. Lei, Y. Zhu, X. Qiu, L. Luo, J. Chen, R. Cheng, Z. Chang, H. Zhang, N.C. Wu, Y. Guo, D. Hou, J. Pei, J. Gao, Y. Hua, Z. Huang, S. Chen, Genomic, transcriptomic, and epigenomic analysis of a medicinal snake, *Bungarus multicinctus*, to provides insights into the origin of Elapidae neurotoxins, Acta Pharm. Sin. B 13 (2023) 2234–2249. 10.1016/j.apsb.2022.11.015.

[41] B. Liu, L. Cui, Z. Deng, Y. Ma, D. Yang, Y. Gong, Y. Xu, T. Lan, S. Yang, S. Huang, The genome assembly and annotation of the many-banded krait, Bungarus multicinctus, GigaByte 2023 (2023) gigabyte82. 10.46471/gigabyte.82.

[42] S. Hurka, K. Brinkrolf, R. Özbek, F. Förster, A. Billion, J. Heep, T. Timm, G. Lochnit, A. Vilcinskas, T. Lüddecke, Venomics of the Central European Myrmicine Ants Myrmica rubra and Myrmica ruginodis, Toxins 14 (2022) 358. 10.3390/toxins14050358.

[43] L. Schulte, M. Damm, I. Avella, L. Uhrig, P. Erkoc, S. Schiffmann, R. Fürst, T. Timm, G. Lochnit, A. Vilcinskas, T. Lüddecke, Venomics of the milos viper (Macrovipera schweizeri) unveils patterns of venom composition and exochemistry across blunt-nosed viper venoms, Front. Mol. Biosci. 10 (2023). 10.3389/fmolb.2023.1254058.

[44] I. Avella, L. Schulte, S. Hurka, M. Damm, J. Eichberg, S. Schiffmann, M. Henke, T. Timm, G. Lochnit, K. Hardes, A. Vilcinskas, T. Lüddecke, Proteogenomics-guided functional venomics resolves the toxin arsenal and activity of *Deinagkistrodon acutus* venom, Int. J. Biol. Macromol. 278 (2024) 135041. 10.1016/j.ijbiomac.2024.135041.

[45] H. Liu, R.G. Sadygov, J.R. Yates, A Model for Random Sampling and Estimation of Relative Protein Abundance in Shotgun Proteomics, Anal. Chem. 76 (2004) 4193–4201. 10.1021/ac0498563.

[46] B. Zybailov, M.K. Coleman, L. Florens, M.P. Washburn, Correlation of Relative Abundance Ratios Derived from Peptide Ion Chromatograms and Spectrum Counting for Quantitative Proteomic Analysis Using Stable Isotope Labeling, Anal. Chem. 77 (2005) 6218–6224. 10.1021/ac050846r.

[47] B. Zybailov, A.L. Mosley, M.E. Sardiu, M.K. Coleman, L. Florens, M.P. Washburn, Statistical Analysis of Membrane Proteome Expression Changes in Saccharomyces cerevisiae, J. Proteome Res. 5 (2006) 2339–2347. 10.1021/pr060161n.

[48] L. Florens, M.J. Carozza, S.K. Swanson, M. Fournier, M.K. Coleman, J.L. Workman, M.P. Washburn, Analyzing chromatin remodeling complexes using shotgun proteomics and normalized spectral abundance factors, Methods 40 (2006) 303–311. 10.1016/j.ymeth.2006.07.028.

[49] M. Damm, M. Karış, D. Petras, A. Nalbantsoy, B. Göçmen, R.D. Süssmuth, Venomics and Peptidomics of Palearctic Vipers: A Clade-Wide Analysis of Seven Taxa of the Genera Vipera, Montivipera, Macrovipera, and Daboia across Türkiye, J. Proteome Res. 23 (2024) 3524–3541. 10.1021/acs.jproteome.4c00171.

[50] M.C. Chambers, B. Maclean, R. Burke, D. Amodei, D.L. Ruderman, S. Neumann, L. Gatto, B. Fischer, B. Pratt, J. Egertson, K. Hoff, D. Kessner, N. Tasman, N. Shulman, B. Frewen, T.A. Baker, M.-Y. Brusniak, C. Paulse, D. Creasy, L. Flashner, K. Kani, C. Moulding, S.L. Seymour, L.M. Nuwaysir, B. Lefebvre, F. Kuhlmann, J. Roark, P. Rainer, S. Detlev, T. Hemenway, A. Huhmer, J. Langridge, B. Connolly, T. Chadick, K. Holly, J. Eckels, E.W. Deutsch, R.L. Moritz, J.E. Katz, D.B. Agus, M. MacCoss, D.L. Tabb, P. Mallick, A cross-platform toolkit for mass spectrometry and proteomics, Nat. Biotechnol. 30 (2012) 918–920. 10.1038/nbt.2377.

[51] A.R. Basharat, Y. Zang, L. Sun, X. Liu, TopFD: A Proteoform Feature Detection Tool for Top-Down Proteomics, Anal. Chem. 95 (2023) 8189–8196. 10.1021/acs.analchem.2c05244.

[52] Q. Kou, L. Xun, X. Liu, TopPIC: a software tool for top-down mass spectrometry-based proteoform identification and characterization, Bioinforma. Oxf. Engl. 32 (2016) 3495– 3497. 10.1093/bioinformatics/btw398.

[53] UniProt Consortium, UniProt: the Universal Protein Knowledgebase in 2025, Nucleic Acids Res. 53 (2025) D609–D617. 10.1093/nar/gkae1010.

[54] B. Buchfink, C. Xie, D.H. Huson, Fast and sensitive protein alignment using DIAMOND, Nat. Methods 12 (2015) 59–60. 10.1038/nmeth.3176.

[55] VenomZone, (n.d.). https://venomzone.expasy.org/ (accessed April 19, 2025).

[56] F. Jungo, A. Bairoch, Tox-Prot, the toxin protein annotation program of the Swiss-Prot protein knowledgebase, Toxicon 45 (2005) 293–301. 10.1016/j.toxicon.2004.10.018.

[57] S. Henikoff, J.G. Henikoff, Amino acid substitution matrices from protein blocks., Proc. Natl. Acad. Sci. 89 (1992) 10915–10919. 10.1073/pnas.89.22.10915.

[58] P.J.A. Cock, T. Antao, J.T. Chang, B.A. Chapman, C.J. Cox, A. Dalke, I. Friedberg, T. Hamelryck, F. Kauff, B. Wilczynski, M.J.L. de Hoon, Biopython: freely available Python tools for computational molecular biology and bioinformatics, Bioinformatics 25 (2009) 1422–1423. 10.1093/bioinformatics/btp163.

[59] F. Teufel, J.J. Almagro Armenteros, A.R. Johansen, M.H. Gíslason, S.I. Pihl, K.D. Tsirigos, O. Winther, S. Brunak, G. von Heijne, H. Nielsen, SignalP 6.0 predicts all five types of signal peptides using protein language models, Nat. Biotechnol. 40 (2022) 1023–1025. 10.1038/s41587-021-01156-3.

[60] P. Jones, D. Binns, H.-Y. Chang, M. Fraser, W. Li, C. McAnulla, H. McWilliam, J. Maslen, A. Mitchell, G. Nuka, S. Pesseat, A.F. Quinn, A. Sangrador-Vegas, M. Scheremetjew, S.-Y. Yong, R. Lopez, S. Hunter, InterProScan 5: genome-scale protein function classification, Bioinformatics 30 (2014) 1236–1240. 10.1093/bioinformatics/btu031.

[61] L. Uhrig, A. Cabrera-Orefice, M. Damm, T. Lüddecke, DATASET - Mass Spectrometry - Snake venom proteogenomics of Bungarus multicinctus from Taiwan, Zenodo (2025). 10.5281/zenodo.12785576.

[62] P. Erkoc, S. Schiffmann, T. Ulshöfer, M. Henke, M. Marner, J. Krämer, R. Predel, T.F. Schäberle, S. Hurka, L. Dersch, A. Vilcinskas, R. Fürst, T. Lüddecke, Determining the pharmacological potential and biological role of linear pseudoscorpion toxins via functional profiling, IScience 27 (2024) 110209. 10.1016/j.isci.2024.110209.

[63] T. Lüddecke, L. Dersch, L. Schulte, S. Hurka, A. Paas, M. Oberpaul, J. Eichberg, K. Hardes, S. Klimpel, A. Vilcinskas, Functional Profiling of the A-Family of Venom Peptides from the Wolf Spider Lycosa shansia, Toxins 15 (2023) 303. 10.3390/toxins15050303.

[64] A. Gomes, P.P. Saha, S. Bhattacharya, S. Ghosh, A. Gomes, Therapeutic potential of krait venom, Toxicon 131 (2017) 48–53. 10.1016/j.toxicon.2017.03.004.

[65] J. Slagboom, C. Kaal, A. Arrahman, F.J. Vonk, G.W. Somsen, J.J. Calvete, W. Wüster, J. Kool, Analytical strategies in venomics, Microchem. J. 175 (2022) 107187. 10.1016/j.microc.2022.107187.

[66] Y.L. Hia, K.Y. Tan, C.H. Tan, Comparative venom proteomics of banded krait (*Bungarus fasciatus*) from five geographical locales: Correlation of venom lethality, immunoreactivity and antivenom neutralization, Acta Trop. 207 (2020) 105460. 10.1016/j.actatropica.2020.105460.

[67] B.C. Offor, B. Muller, L.A. Piater, A Review of the Proteomic Profiling of African Viperidae and Elapidae Snake Venoms and Their Antivenom Neutralisation, Toxins 14 (2022) 723. 10.3390/toxins14110723.

[68] L.-L. Shan, J.-F. Gao, Y.-X. Zhang, S.-S. Shen, Y. He, J. Wang, X.-M. Ma, X. Ji, Proteomic characterization and comparison of venoms from two elapid snakes (*Bungarus multicinctus* and *Naja atra*) from China, J. Proteomics 138 (2016) 83–94. 10.1016/j.jprot.2016.02.028.

[69] U. Rashmi, S. Bhatia, M. Nayak, S. Khochare, K. Sunagar, Elusive elapids: biogeographic venom variation in Indian kraits and its repercussion on snakebite therapy, Front. Pharmacol. 15 (2024). 10.3389/fphar.2024.1443073.

[70] K. Sunagar, S. Khochare, R.R. Senji Laxme, S. Attarde, P. Dam, V. Suranse, A. Khaire, G. Martin, A. Captain, A Wolf in Another Wolf’s Clothing: Post-Genomic Regulation Dictates Venom Profiles of Medically-Important Cryptic Kraits in India, Toxins 13 (2021) 69. 10.3390/toxins13010069.

[71] F.L. Yuan, T.-L. Prigge, Y.-H. Sung, C. Dingle, T.C. Bonebrake, Two Genetically Distinct yet Morphologically Indistinct Bungarus Species (Squamata, Elapidae) in Hong Kong, Curr. Herpetol. 41 (2022) 114–124. 10.5358/hsj.41.114.

[72] Z.-N. Chen, S.-C. Shi, G. Vogel, L. Ding, J.-S. Shi, Multiple lines of evidence reveal a new species of Krait (Squamata, Elapidae, Bungarus) from Southwestern China and Northern Myanmar, ZooKeys 1025 (2021) 35–71. 10.3897/zookeys.1025.62305.

[73] A.E. Levitón, G.O.U. Wogan, M.S. Koo, G.R. Zug, R.S. Lucas, J.V. Vindum, The Dangerously Venomous Snakes of Myanmar, Proc. Calif. Acad. Sci. 54 (2003).

[74] J. Harris, A. Goonetilleke, ANIMAL POISONS AND THE NERVOUS SYSTEM: WHAT THE NEUROLOGIST NEEDS TO KNOW, J. Neurol. Neurosurg. Psychiatry 75 (2004) iii40–iii46. 10.1136/jnnp.2004.045724.

[75] S. Prasarnpun, J. Walsh, S.S. Awad, J.B. Harris, Envenoming bites by kraits: the biological basis of treatment-resistant neuromuscular paralysis, Brain 128 (2005) 2987– 2996. 10.1093/brain/awh642.

[76] A. Silva, K. Maduwage, M. Sedgwick, S. Pilapitiya, P. Weerawansa, N.J. Dahanayaka, N.A. Buckley, C. Johnston, S. Siribaddana, G.K. Isbister, Neuromuscular Effects of Common Krait (Bungarus caeruleus) Envenoming in Sri Lanka, PLoS Negl. Trop. Dis. 10 (2016) e0004368. 10.1371/journal.pntd.0004368.

[77] R.W. Dixon, J.B. Harris, Nerve terminal damage by beta-bungarotoxin: its clinical significance, Am. J. Pathol. 154 (1999) 447–455. 10.1016/s0002-9440(10)65291-1.

[78] C.C. Chu, S.T. Chu, S.W. Chen, Y.H. Chen, The non-phospholipase A2 subunit of beta-bungarotoxin plays an important role in the phospholipase A2-independent neurotoxic effect: characterization of three isotoxins with a common phospholipase A2 subunit, Biochem. J. 303 (Pt 1) (1994) 171–176. 10.1042/bj3030171.

[79] R. Doley, R.M. Kini, Protein complexes in snake venom, Cell. Mol. Life Sci. CMLS 66 (2009) 2851–2871. 10.1007/s00018-009-0050-2.

[80] Y. Tian, Z. Liu, L. Ma, Y. Yu, Q. Shi, S. Zhao, Y. Zhou, Forensic identification of a fatal snakebite from Bungarus multicinctus (Chinese krait) by pathological and toxicological findings: a case report, Forensic Sci. Med. Pathol. 18 (2022) 497–502. 10.1007/s12024-022-00517-x.

[81] M. Gao, X. Zhang, T. Jian, C. Sun, G. Yu, Y. Gao, B. Kan, X. Jian, Medical management of a child treated for two unique envenomation episodes via captive snakes in a 60-day period: A case report, Heliyon 10 (2024). 10.1016/j.heliyon.2024.e40245.

[82] S.P. Mackessy, ed., Handbook of Venoms and Toxins of Reptiles, 2nd ed., CRC Press, Boca Raton, 2021. 10.1201/9780429054204.

[83] J.W. Fox, S.M.T. Serrano, Structural considerations of the snake venom metalloproteinases, key members of the M12 reprolysin family of metalloproteinases, Toxicon 45 (2005) 969–985. 10.1016/j.toxicon.2005.02.012.

[84] R.M. Kini, C.Y. Koh, Metalloproteases Affecting Blood Coagulation, Fibrinolysis and Platelet Aggregation from Snake Venoms: Definition and Nomenclature of Interaction Sites, Toxins 8 (2016) 284. 10.3390/toxins8100284.

[85] E.G. Rowan, What does β-bungarotoxin do at the neuromuscular junction?, Toxicon 39 (2001) 107–118. 10.1016/S0041-0101(00)00159-8.

